# Cryo-electron tomography of eel sperm flagella reveals a molecular “minimum system” for motile cilia

**DOI:** 10.1101/2023.07.16.549168

**Authors:** Jason R. Schrad, Gang Fu, Whitney E. Hable, Alexandra M. Tayar, Kenneth Oliveira, Daniela Nicastro

## Abstract

Cilia and flagella play a crucial role in the development and function of most eukaryotic organisms. The structural core of the flagellum, the axoneme, is conserved in most eukaryotes and careful regulation of dynein motors within the axoneme is required for proper ciliary beating. The sperm flagellum from the American eel (*Anguilla rostrata*) has been shown to lack many of the canonical axonemal proteins, including the radial spokes, the central pair complex, and possibly even the outer row of dynein arms, presenting a “minimal” flagellar system. Here, we present cryo-electron tomography analysis of the eel sperm flagellum. We identified two states for the eel sperm flagellum within our tomograms, narrow and wide, and found that the flagellum started narrow near the sperm head and widened distally. Subtomogram averages revealed that the eel sperm flagellum has retained remnants of the missing regulatory complexes, including a short radial spoke 3 complex, basal components of radial spokes 1 and 2, and an outer dynein arm docking complex. We also describe unique structural features of the *A. rostrata* sperm flagellum, such as a unique pattern of holes at the inner junction and an accessory complex located at the “outer” junction. Finally, we discuss the consequences of losing key regulatory factors for the eel sperm flagellum and hypothesize several evolutionary factors that may have led to their loss. Together, our results shed light onto the structure and function of the eel sperm axoneme and provide insight into the minimum requirements for proper ciliary beating.

## Introduction

Cilia (flagella) are hair-like appendages that extend from the surface of many eukaryotic cells (Satir and Christensen, 2007; Takahashi, 1984). These organelles are conserved amongst eukaryotes, including most protists (*e.g.,* the green algae *Chlamydomonas* and *Tetrahymena*) and metazoa (*e.g., Drosophila*, mice, and humans) (Satir and Christensen, 2007; Takahashi, 1984). Cilia are responsible for a wide range of biological functions, including cellular locomotion, signal transduction, developmental regulation, and fluid clearance (May-Simera et al., 2017; Moran et al., 1977; Pan and Snell, 2007; Satir and Christensen, 2007; Viswanadha et al., 2017). These organelles are generally split into two categories: primary cilia and motile cilia (Satir and Christensen, 2007).

Primary cilia, also known as immotile cilia, largely function in a regulatory and/or signal transduction manner within cells and organisms (May-Simera et al., 2017; Moran et al., 1977; Pan and Snell, 2007). For example, primary cilia within *C. elegans* sensory neurons are responsible for mechanosensation (Doroquez et al., 2014; Nechipurenko et al., 2017). Motile cilia, on the other hand, are cilia that are capable of movement. These cilia are primarily used to propel locomotion in many single celled organisms or to move fluids across tissues in multicellular organisms (Satir and Christensen, 2007; Takahashi, 1984). Motile cilia also play a role in several stages of vertebrate development, including the development of the heart and the left-right body axis (Djenoune et al., 2022; Nonaka et al., 1998; Willaredt et al., 2012). Defects in either primary or motile cilia can lead to severe defects in cells and organisms. Mutations in just one of the over 600 proteins in a motile cilium can result in severe motility defects or paralysis (Bower et al., 2018; Urbanska et al., 2018; Urbanska et al., 2015). In humans, ciliary defects can lead to numerous diseases - called ciliopathies - including primary ciliary dyskinesia (Hjeij et al., 2014; Wirschell et al., 2011; Yamamoto et al., 2013), *situs inversus* (Cannarella et al., 2020; Matwijiw et al., 1987), male infertility (Coutton et al., 2018), and polycystic kidney disease (Huang and Lipschutz, 2014). Although ciliary beating plays a critical role in eukaryotic life cycles, the structures and molecular mechanisms that control beating are not yet fully understood.

Cilia are comprised of a highly periodic structural backbone, known as the axoneme, which usually consists of a circular array of nine doublet microtubules (DMTs) (Fig. 1A) (Sale and Satir, 1977; Witman et al., 1978). Most motile cilia have a [9+2] microtubule arrangement, meaning nine DMTs surrounding a pair of interconnected singlet microtubules, called the central pair complex (CPC). In contrast, most primary cilia contain [9+0] axonemes, lacking a CPC (Hoey et al., 2012; Satir and Christensen, 2007). Although most motile cilia contain [9+2] axonemes, there are some notable exceptions, including the [6+0] motile cilium of the protist *Lecudina tuzetae* (Schrevel and Besse, 1975), [9+0] motile nodal cilia (Sulik et al., 1994), and [10+0], [16+0], or [20+0] motile sperm flagella found in some insect species (Dallai et al., 2006).

**Figure 1:**
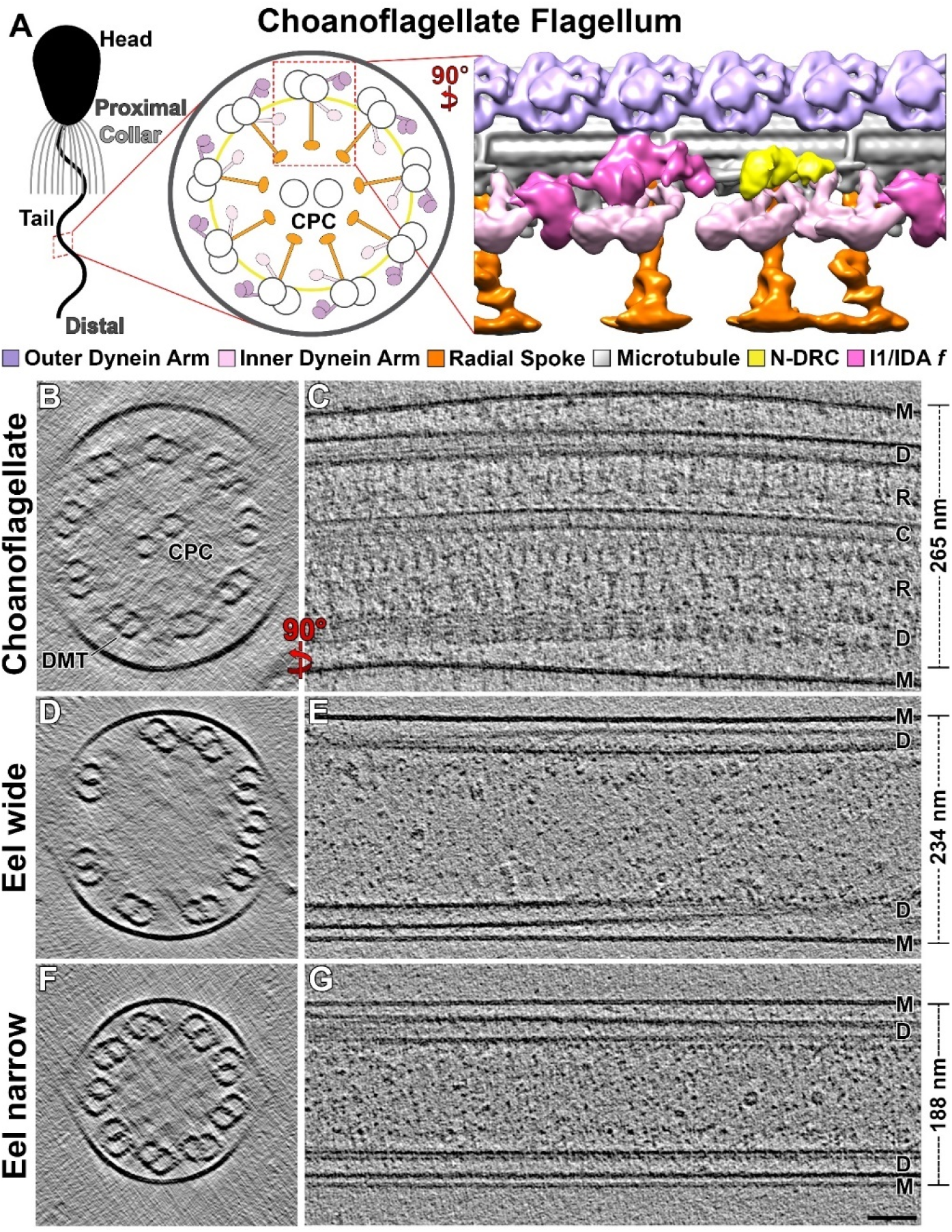
Cryo-ET provides a comparison of [9+2] and [9+0] motile flagella structures. A) Schematic representation of the flagellar organization and 96-nm axonemal repeat unit (A) of the [9+2] motile flagellum of the choanoflagellate *Salpingoeca rosetta*. *S. rosetta* cells utilize a specialized “collar” structure that surrounds the distal region of the flagellum. B-C) Cross sectional (B) and longitudinal (C) tomographic slices of a representative *S. rosetta* flagellum. D-G) Cross sectional (D, F) and longitudinal (E, G) tomographic slices of American eel (*Anguilla rostrata*) flagella. Two structural classes of eel flagella were observed, “wide” (D-E) or “narrow” (F-G). Labels at the right of panels C, E, and G highlight the membrane (M), DMTs (D), radial spokes (R), and CPC (C). The scale bar in G represents 50 nm and is valid for panels B-G. For all longitudinal slices and isosurface renderings, the proximal end of the repeat is on the left, unless otherwise noted. For all cross-sectional slices and renderings, the proximal end of the repeat faces the viewer, unless otherwise noted.

Despite the variations in the number of microtubules and biological functions, the periodic axoneme structure itself is highly conserved (Linck et al., 2014; Nicastro et al., 2011). The base structural subunit of the axoneme is a cassette of structural features that can be found every 96 nm along the length of each DMT (Fig. 1A) (Nicastro et al., 2011; Nicastro et al., 2006). This repeating subunit consists of the DMT and numerous proteins that stabilize the DMT (*e.g.,* microtubule inner proteins or MIPs), drive flagellar beating (dynein motors), or transduce signals to control beating (*e.g.,* the nexin-dynein regulatory complex or the radial spokes). In a [9+2] axoneme, each DMT is connected to the CPC via two or three radial spoke complexes (RSs) per 96-nm repeat unit (Fig. 1A). Several connections link neighboring DMTs, including dynein arms and the nexin-dynein regulatory complex (N-DRC) (Heuser et al., 2009). DMTs are decorated with two rows of dyneins, the inner and outer dynein arms (IDAs and ODAs, respectively), that are attached to the A tubule of one DMT and reach towards the B tubule of the adjacent DMT. Activation of the dynein motor domains drives sliding between neighboring DMTs. Inter-DMT linkers, *e.g.,* such as the N-DRC, restrict this sliding movement, causing the cilia to bend (Heuser et al., 2009; Lin and Nicastro, 2018).

For motile cilia to function effectively, their cilia must beat properly, with tight control of beat frequency, bend direction and amplitude, and beat propagation. To achieve proper ciliary beating, organisms must carefully regulate dynein activation through axonemal signal transduction pathways (Lin and Nicastro, 2018; Viswanadha et al., 2017; Wirschell et al., 2011). Signals begin at the CPC, propagate up the RSs to the IDAs, and then on to the ODAs via a specialized ODA-IDA linker region (Gui et al., 2019; Oda et al., 2013). This signal transduction pathway is critical for proper flagellar beating. Loss of, or defects in, components of the axoneme can result in cilia that are paralyzed or have impaired motility. For example, the RS-less *pf14* mutant (Diener et al., 1993) and the CPC-less *pf18* mutant in *Chlamydomonas* (Adams et al., 1981) have paralyzed flagella or flagella with impaired motility, respectively. Previous studies have shown, however, that there are naturally occurring motile cilia that lack critical components of the CPC-RS-dynein pathway. For example, *Chlamydomonas* flagella have a short third radial spoke (Barber et al., 2012; Heuser et al., 2012). Also, motile nodal cilia, which are thought to play a role in determining the left-right body axis during development, (Buceta et al., 2005; Nonaka et al., 1998; Sulik et al., 1994), and sperm cells from the Asian horseshoe crab (Ishijima et al., 1988) both utilize [9+0] axonemes within their motile cilia. It is not clear how motile [9+0] flagella regulate their dynein activity, and thus their ciliary beating, without the canonical CPC-RS-dynein signaling pathways.

In this study, we used the [9+0] sperm flagellum from the American eel (*Anguilla rostrata*) as a model system to investigate ciliary motility. Similar to the horseshoe crab, classical TEM studies have shown that sperm cells from Anguillid eels (*Anguilla spp.*) lack RSs and a CPC *(Woolley, 1997; Woolley, 1998b)*. Unlike the horseshoe crab (Ishijima et al., 1988), the eel flagellum lacks ODAs, leaving only the N-DRC and IDAs as inter-DMT connections. Without ODAs, the *A. rostrata* flagellum represents the simplest known motile cilium, providing a potential “minimum ensemble” of axonemal structures that is sufficient for motility. To investigate this ensemble, we analyzed the 3D structure of the eel sperm axoneme using cryo-electron tomography (cryo-ET) and subtomogram averaging. We characterized the 96-nm repeating subunit of the eel sperm axoneme and compared it to the ciliated choanoflagellate *Salpingoeca rosetta*. Through this comparison, we discovered that the structure of the eel sperm axoneme has retained the overall architecture of the canonical axoneme but also has several inconsistencies. We also investigated the eel sperm flagellum proteome and the genome of the closely related European eel (*Anguilla anguilla*) to identify known axonemal genes/proteins in the eel flagellum. From these combined analyses, we generated evolutionary hypotheses on the simplicity of the eel flagellum and provide an insight into the structures required for ciliary beating in its purest form.

## Results

### The eel sperm flagellum has a variable width

Although the ultrastructure of the eel sperm axoneme has been described previously (Woolley, 1997), the molecular composition and organization within the flagellum remains unstudied. To generate structural information on the eel sperm tail, we carried out cryo-ET on intact, vitrified sperm cells. Sperm cells were collected from mature male eels and were rapidly frozen for cryo-ET imaging. Initial tomographic reconstructions support Woolley’s conclusions that the eel sperm flagellum lacks RSs and the CPC (Fig. 1D-G) (Woolley, 1997). We then compared tomograms of the eel sperm flagellum with a representative tomogram of the [9+2] flagellum of the choanoflagellate *Salpingoeca rosetta* (Fig. 1B-C) (Pinskey et al., 2022). *S. rosetta* is a ciliated, single-celled organism that is among the closest unicellular relatives of animals. Similar to sperm cells, the choanoflagellate cells are equipped with a single flagellum, but unlike sperm, the *S. rosetta* flagellum has been modified to aid in predation. These modifications include a “collar” of microvilli surrounding the proximal region of the flagellum (Fig. 1A) as well as a matrix of vane filaments extending from the flagellar membrane (Hibberd, 1975). Despite these external differences, the *S. rosetta* axoneme adopts a [9+2] architecture and the axoneme repeat unit resembles that of other eukaryotic flagella (Pinskey et al., 2022).

On average, the eel flagellum was smaller in diameter (187 nm) than that of *S. rosetta* (265 nm). At an average diameter of 187 nm, the eel flagellum is wider than may be expected by simply removing the CPC and RSs from the axoneme of another species. When taken together, the CPC (∼70 nm wide) and RSs (∼40 nm long, x2) occupy the innermost 150 nm of the choanoflagellate flagellum (Pinskey et al., 2022). Simply removing this 150 nm, one might expect to be left with a flagellum that is ∼100 nm wide, although the inter-DMT spacing in this hypothetical axoneme would be incredibly small. It is possible that the smaller eel flagella represent the minimum inter-DMT spacing required for ciliary beating.

The flagella depicted in our tomograms varied in diameter (Fig. 1D-G), ranging from 166-225 nm. For traditional [9+2] flagella, an increased diameter typically corresponds to structural disturbances in the axoneme (*i.e.,* axoneme compression during freezing). Some of the wider eel flagella were compressed, but many retained a circular axoneme (Fig. 1D-E, Fig. 2). Severely compressed flagella were not included in subsequent subtomogram averaging or analysis. Flagella were separated into “narrow” and “wide” categories based on their diameter. Representative wide and narrow flagella can be seen in Fig. 1D-E and 1F-G, respectively. DMTs in the narrow flagella were regularly spaced around the flagellar cross-section (Fig. 1F) whereas wide flagella had variable inter-DMT spacings (Fig. 1D). The spacing between DMTs in the narrow flagella (and those in well-ordered sections of the wide flagella) is smaller than the spacing seen in other organisms (see Discussion).

**Figure 2:**
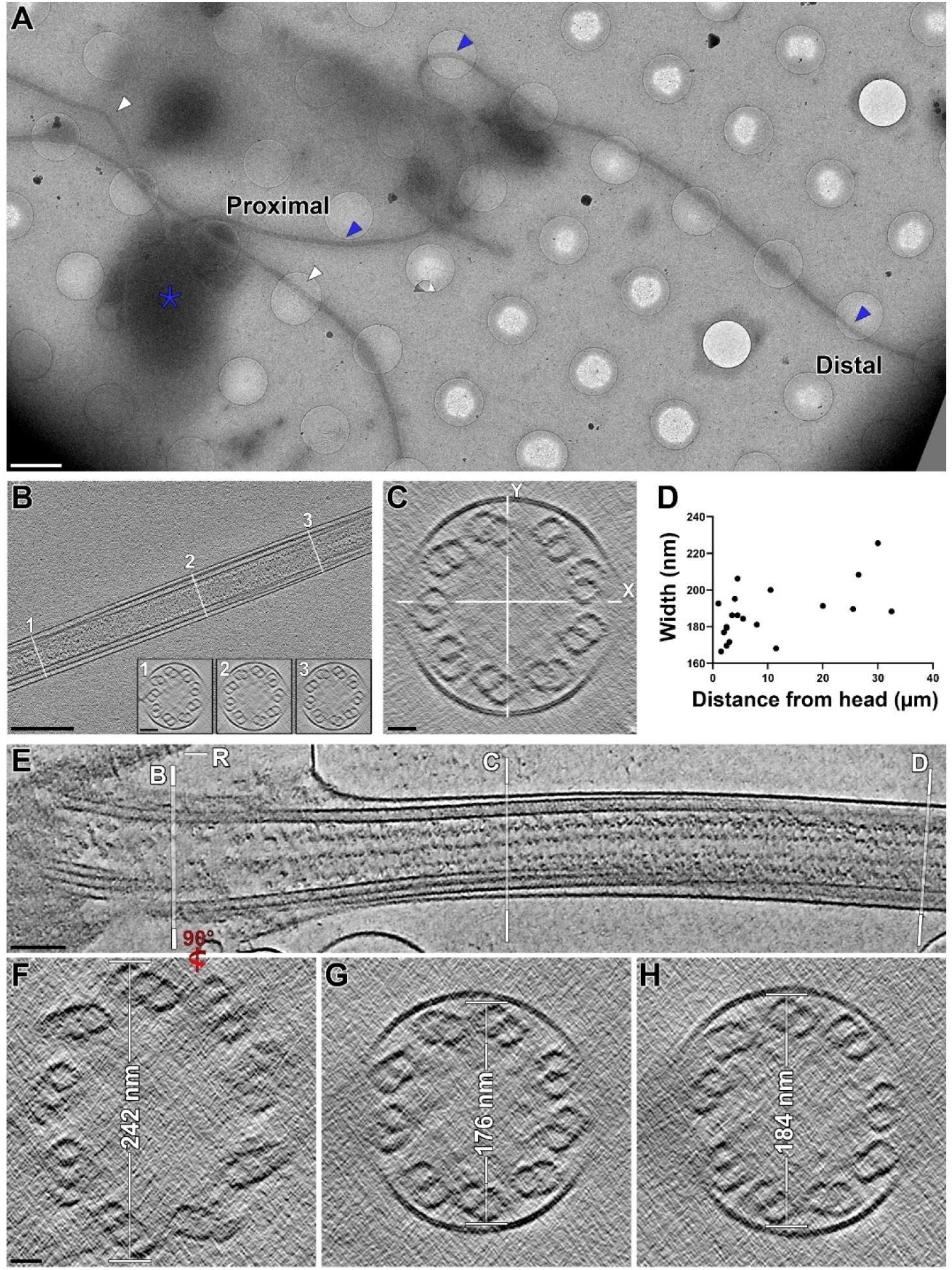
The eel flagellum widens distally along the flagellum but the axoneme structure is retained. A) Cryo-EM map of a vitrified eel sperm cell with a visible sperm head (blue star) and tail (blue arrowheads). Note that the flagella from other sperm cells are also visible (white arrowheads). B) Tomographic slice of an eel sperm flagellum. White lines correspond to the three locations along the flagellum where width measurements were taken. Cross sectional slices at each location are shown in the corresponding inset image. C) Cross sectional tomographic slice depicting an uncompressed (round) flagellum. White lines provide examples of width measurements. D) Scatter plot depicting the width of the flagella used in these averages compared to the distance along the sperm tail that the tilt series were collected at. Note the cluster of narrow flagella measured within the first few microns of the flagellum. E) Longitudinal tomographic slice of the base and proximal region of an eel sperm flagellum. Note the “striated rootlet” (R) extending from the sperm head, visible in the top-left portion of this panel. F) Cross sectional tomographic slice of the basal region of an eel flagellum. Note the triplet microtubules corresponding to the basal body. G-H) Cross sectional tomographic slices of two proximal regions of an eel flagellum. Note the increased width of the flagellum and the increased disorder of the DMT arrangement in the more distal segment (H). Scale bar in panel A represents 100 nm. Scale bar in panel B represents 50 nm and is valid for panels B-C. White lines represent the locations of views represented in subsequent panels. G-H) Isosurface renderings of the eel 96-nm repeat unit average in cross sectional (G) or longitudinal (H) orientation. A key describing the correlation between colors and axonemal features can be found below the isosurface rendering. Scale bars represent 2 µm in panel A, 250 nm in B (50 nm in insets), 100 nm in panel E, and 25 nm in panels C and F (valid for F-H).

Differences in the axoneme structure within spatially regulated regions of the flagellum (*i.e.,* proximal-distal differences) have been demonstrated previously (Lin et al., 2012) and some of these differences are thought to play a role in the regulation of ciliary beating (Lin and Nicastro, 2014; Lin and Nicastro, 2018). To investigate the potential proximal-distal relationship of the width of the eel sperm flagellum, we first determined the location of each tilt series along the flagellum. Using grid square-level maps (Fig. 2A), we identified the point along the flagellum where each tilt series was collected and measured the distance along the flagellum from these points to the sperm head. We then measured the width of each flagellum at three points throughout the tomogram (Fig. 2B-C). We compared the average flagellar width to the distance along the flagellum and found that flagella that we classified as “narrow” tended to be more proximal (average distance of 6.5 µm from the head) than “wide” flagella (18 µm from the head) (Fig. 2D). Differences in axoneme diameter can also be seen within a single flagellum (Fig. 2E-H). For example, along the basal and proximal regions of one eel sperm flagellum (Fig. 2E), the axoneme begins at the basal body, transitions to a small axoneme (Fig. 2G), and widens at the distal end of the visible region (Fig. 2H). This tomogram also includes information on both the basal body and transition zone of the eel flagellum (Fig. 2E-F). While the basal body does consist of the traditional triple microtubules, the eel sperm flagellum appears to lack many of the traditional accessory structures associated with the basal body and transition zone, including the ciliary necklace and the Y-fibers (Awata et al., 2014; Gilula and Satir, 1972). This tomogram also contains a “striated rootlet” structure (“R” in Fig. 2 E) previously observed in both Atlantic and Pacific eel species (Okamura et al., 2000; Todd, 1976).

To determine if there were structural differences between the narrow and wide flagella, we averaged 96-nm repeat units from each flagella category separately (Fig. 3A-D). Density corresponding to the neighboring DMT (DMT_n+1_) is much weaker in the wide average (Fig. 3A) but obviously visible in the cross section of the narrow average (Fig. 3C). The relatively strong density in the narrow flagella suggests that the inter-DMT spacing in these flagella is more regular than in the wide flagella. This spacing disparity can be directly observed by comparing Fig. 1D and Fig. 1F. Within the DMT of interest (DMT_n_), there appear to be no structural differences between the wide and narrow flagella averages (Fig. 3A-D), therefore, we combined 96-nm repeating subunits from all flagella in future experiments (Fig. 3E-H).

**Figure 3:**
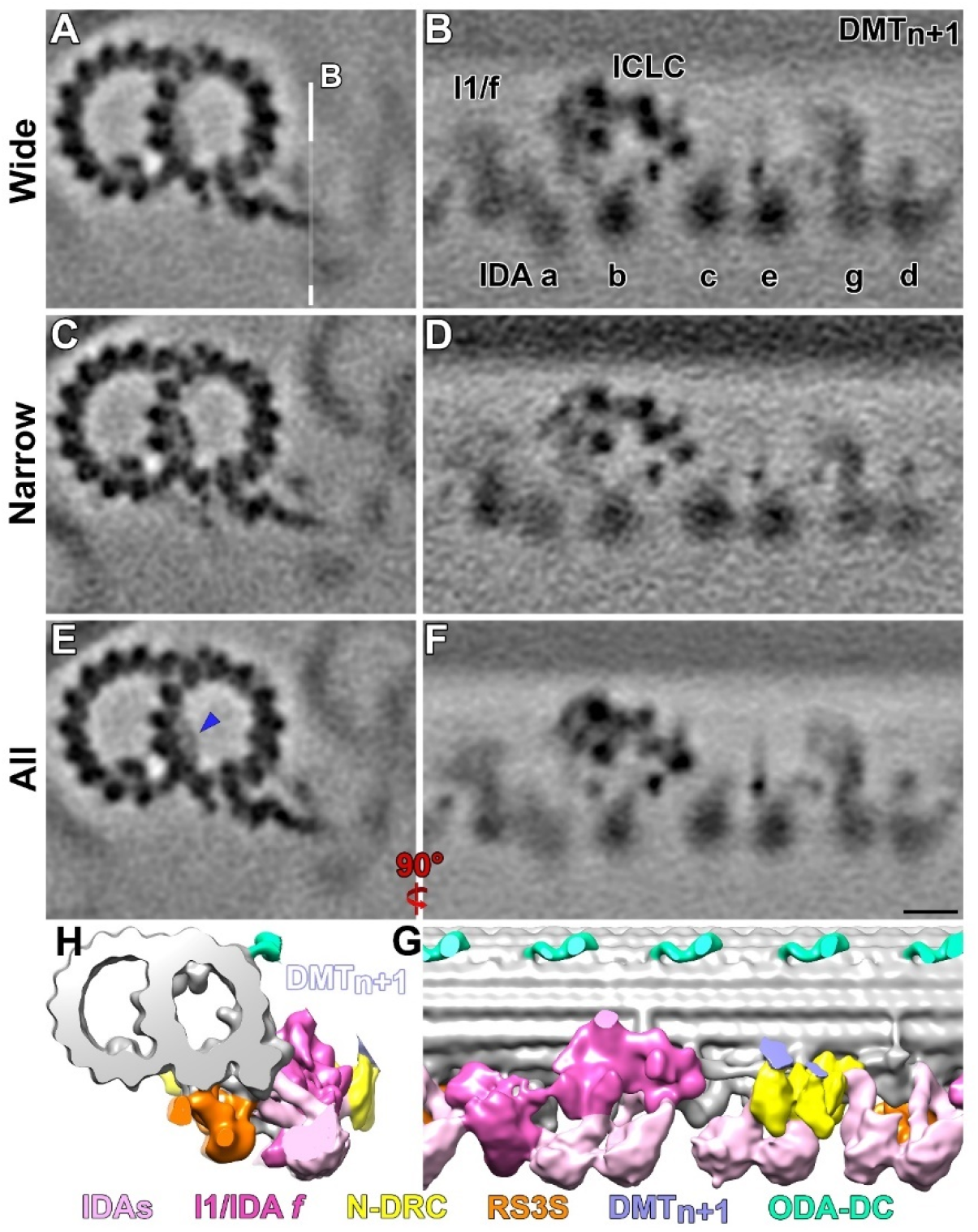
Small and wide regions of the eel sperm flagellum are structurally similar. A-D) Cross sectional (A, C) and longitudinal (B, D) slices of subtomogram averages generated from the wide (A-B) or narrow (C-D) eel sperm flagella. Note the structural similarity between the two averages. Key structural features, including the IDAs, the ICLC complex, and the neighboring DMT are highlighted (black labels). There were no major structural differences between the narrow and wide averages, so the particles were combined and averaged. E-F) Cross sectional (E) and longitudinal (F) slices of the combined subtomogram average. Note the decrease in the density of the neighboring DMT and the MIP4 region (blue arrowhead) in the combined averaged compared to the narrow average. G-H) Isosurface renderings of the eel 96-nm repeat unit average in cross sectional (G) or longitudinal (H) orientation. To aid in visualizing the ODA-DC structure, density for the neighboring DMT was computationally removed using Chimera’s Volume Eraser functionality. A key describing the correlation between colors and axonemal features can be found below the isosurface rendering. The scale bar in panel F represents 10 nm and is valid for panels A-F. White lines represent the locations of views represented in subsequent panels.

### The eel sperm flagellum has retained remnants of canonical regulators of flagellar motility

As demonstrated through traditional thin section electron microscopy (Woolley, 1997) and supported by our tomograms, the eel sperm flagellum lacks many of the structures thought to regulate flagellar beating. Among these missing structures are the CPC, the RSs, and the ODAs. Through subtomogram averaging and proteomic analyses, we have found remnants of the RSs and ODAs within the eel axoneme. CPC remnants were not identified through either of these analyses. The eel flagellum does not utilize the traditional sinusoidal ciliary beating pattern employed by many [9+2] flagella (Lin and Nicastro, 2018; Woolley, 1998a; Woolley, 1998b). Instead, the eel flagellum has a helical beating pattern (Movie S1) (Woolley, 1998a; Woolley, 1998b). It is likely that the lack of canonical regulation structures (*e.g.,* CPC or RSs) leads to this atypical beating pattern.

Consistent with previous conventional EM studies (Woolley, 1997), our tomograms demonstrated that the eel flagellum lacks full-length RSs (Figs. 1-3). Subtomogram averaging of the eel 96-nm axoneme repeat revealed a 14-nm long structure that connects to the A-tubule and hangs below the IDAs (Fig. 3H, Fig. 4C, E, G). This structure is located distally from the N-DRC, between IDA *d* and IDA *g* (Fig. 4C), matching the location of RS3 in organisms that utilize three full-length RSs (Lin and Nicastro, 2018). The shape of this structure is reminiscent of the RS3 short (RS3S) structure found in *Chlamydomonas* (Barber et al., 2012; Heuser et al., 2012). *Chlamydomonas* utilizes two full-length RSs (RS1 & RS2) and a third, truncated RS *in lieu* of a full-length RS3.

**Figure 4:**
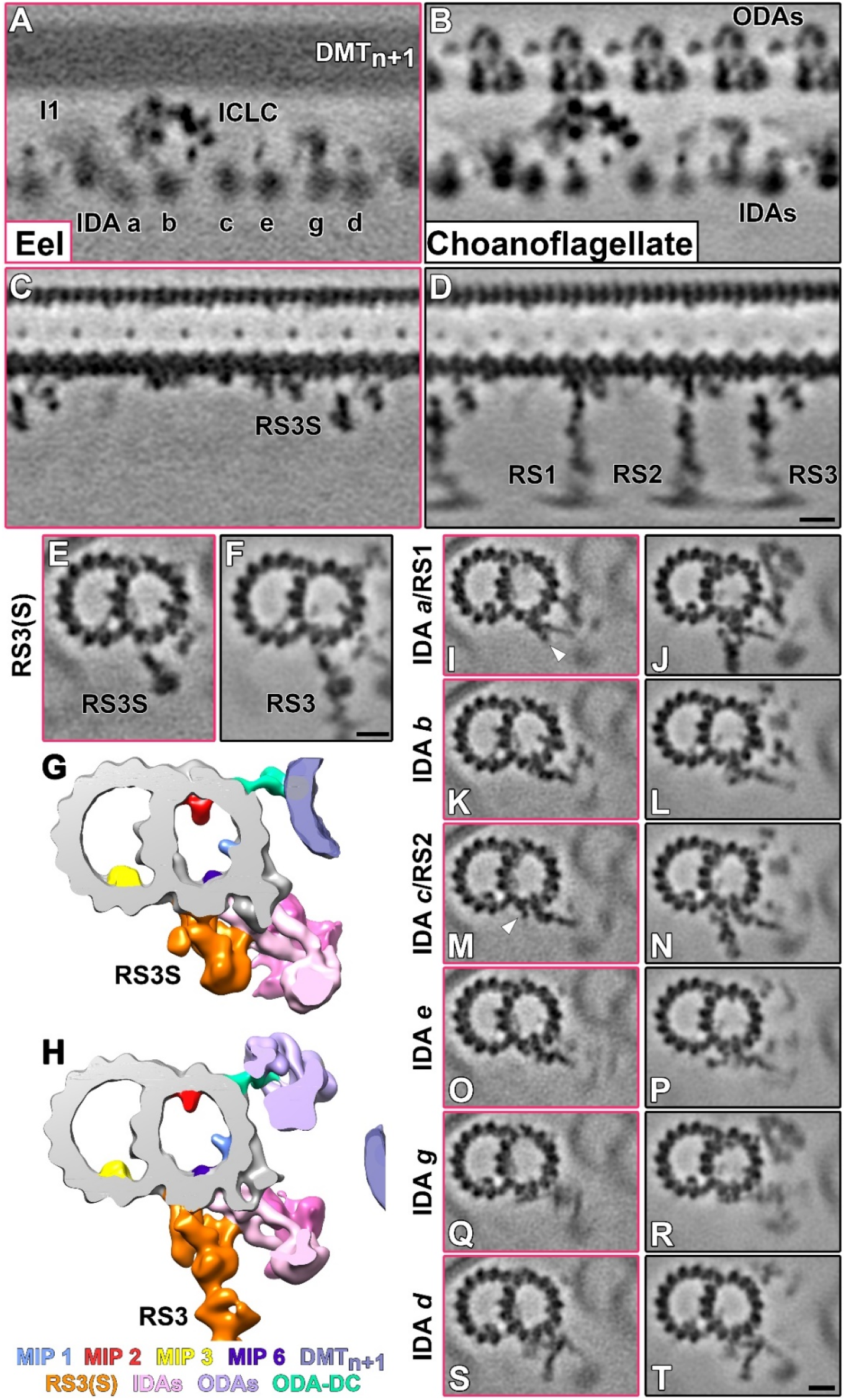
The eel sperm flagellum demonstrates conserved IDA docking and retains radial spoke remnants. A-D) Longitudinal slices of the averaged 96-nm repeat units from the eel sperm flagellum (A, C) and the *S. rosetta* flagellum (B, D). Slices in panels A & B are focused on the IDA dynein heads (black labels) and on the RSs in panels C & D. Tomographic slices are outlined corresponding to the source organism: eel – magenta, *S. rosetta* – black. E-H) Cross sectional tomographic slices (E-F) and isosurface renderings (G-H) of the eel (E, G) and *S. rosetta* (F, H) axonemal repeat units, centered on the location of RS3(S). The eel sperm flagellum contains an RS3S structure, similar to that seen in *Chlamydomonas*. Note the similarity in the eel RS3S structure (orange) and the basal region of the *S. rosetta* RS3. Colors in the isosurface renderings highlight conserved structural features and can be identified using the legend. I-T) Cross sectional tomographic slices depicting the indicated IDA tail and/or RS. Structures resembling components of RS1 and RS2 are also present along the eel sperm flagellum (cyan and green arrowheads, respectively). Scale bars represent 10 nm. The scale bar in panel D is valid for A-D, in panel F for E-F, and in panel T for I-T.

Much of the previous work towards identifying RS proteins has been carried out in *Chlamydomonas*, and the component proteins of RS1/2 are known (Barber et al., 2012; Gui et al., 2021b; Huang et al., 1981; Yang et al., 2006). Identification of the protein components of RS3 has proven more elusive. In *Chlamydomonas*, however, the RS3S complex has been shown to associate with several proteins, including those in the calmodulin and spoke-associated complex (CSC) (Dymek et al., 2011; Heuser et al., 2012; Urbanska et al., 2015). To investigate the presence of RS- and CSC-associated proteins in the eel flagellum, we performed mass spectroscopy on isolated eel sperm flagella. This proteomic investigation produced hits for several RS/CSC proteins, including calmodulin, the RS chaperone DnaJ (HSP40), as well as two proteins (FAP206 and MORN3) that are involved in RS2 docking onto the microtubule (Table 1).

**Table 1:**
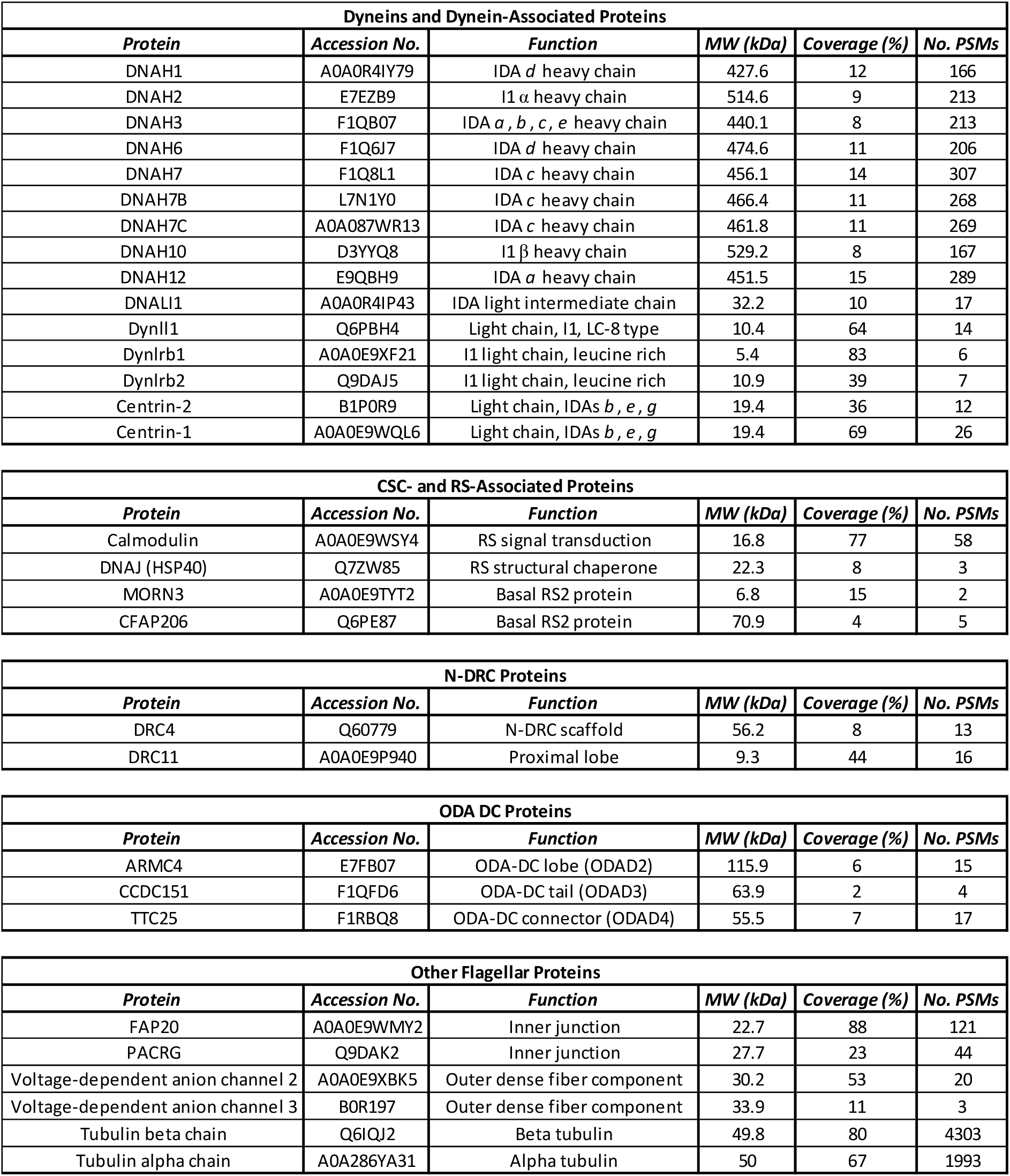
Axonemal proteins identified through mass spectrometry of eel sperm flagella. Proteins with less than 5% coverage and fewer than three peptide spectral matches (PSMs) were excluded from this list. Dynein proteins are labeled according to the unified nomenclature laid out in (Braschi et al., 2022).

While there are not RS3S-sized structures at the locations of RS1 or RS2, there appear to be remnants of each of these other RSs present in the eel axoneme and/or proteome (Fig. 4C-D). In RS-containing organisms, several of the IDAs dock to the microtubule through interactions with RS proteins. IDA *c* binds to the front spoke (FAP206) of RS2 in *Tetrahymena* (Vasudevan et al., 2015) and IDA *b* is thought to interact with FAP253 in *Chlamydomonas* (Gui et al., 2021b). FAP206 was present in the eel mass spectrometry results (Table 1). Structural remnants of several RS1- and RS2-associated proteins are present in the eel axoneme (Fig. 4I-J, M-N). Electron density like the front spoke-IDA *c* region in the *S. rosetta* axoneme (Fig. 4N) can be seen where IDA *c* docks to the A-tubule of the eel DMT. The axonemal ruler (a complex of Ccdc39/Ccdc40) runs along the length of the axoneme repeat unit between protofilaments A2 and A3 (white arrowhead in Fig. 5A). Finally, where the IDA *a* tail interacts with the DMT in the eel axoneme, a density spans the A2-A3 space and hangs down from the eel microtubule (Fig. 4I-J). This structure is reminiscent of the density occupied by FAP207 (MORN repeat containing protein 3, MORN3) in the *Chlamydomonas* RS1 (Gui et al., 2021b). It remains unclear if these supposed remnants play a role in the function of the eel axoneme, but their presence in an RS-less organism suggests that they could be critical for IDA docking (below).

**Figure 5:**
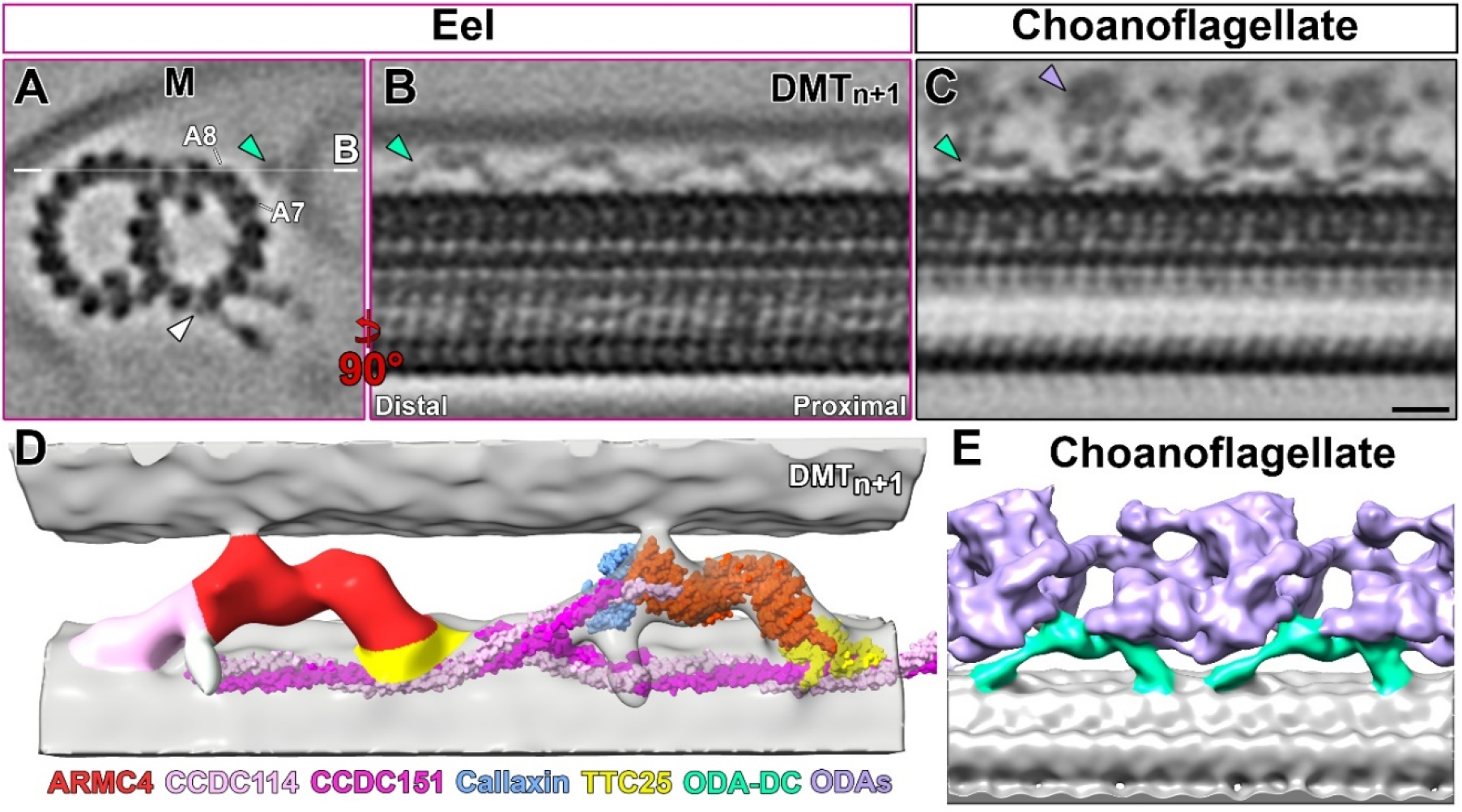
The eel sperm flagellum has retained an ODA docking complex (ODA-DC). A) Cross-sectional slice of the eel sperm flagellum 96-nm repeat unit. Note the relative proximity of the flagellar membrane (M) and the neighboring DMT in the average. A structure is present between protofilaments A7 and A8 (cyan arrowhead). This structure is reminiscent of an ODA docking complex (ODA-DC). The white line in A depicts the location and orientation of the slice in B. The white arrowhead highlights the axonemal ruler. B-C) Longitudinal slices of the eel sperm (B) and *S. rosetta* (Choanoflagellate, C) ODA-DCs. Panels B and C are oriented with the distal region to the left and proximal to the right. The eel ODA-DC appears structurally similar to that of Choanoflagellates (Opisthokonta). D-D) Isosurface renderings of the eel sperm (D) and *S. rosetta* (E) ODA-DCs. The metazoan (Choanoflagellate) ODA-DC consists of ARMC4 (red), calaxin (blue), TTC25 (yellow), and the CCDC114/CCDC151 complex (light pink/magenta). The cryo-EM structure of the bovine ODA-DC (PDB 7RRO) docks into the eel ODA-DC density, with the exception of calaxin. There is a protrusion on the opposite side of the eel ARMC4 density (towards the viewer) that may be the location of the eel calaxin subunit. The scale bar in C represents 10 nm and is valid for A-C.

### The eel flagellum lacks ODAs but has retained an ODA docking complex (ODA-DC)

Previous conventional EM experiments suggested that the eel sperm flagellum lacked the outer row of dynein arms (Woolley, 1997; Woolley, 1998b). Cross sections of eel sperm flagella cryo-tomograms corroborate the lack of ODAs (Fig. 1D/F). Subtomogram averaging of the eel 96-nm axoneme repeat unit revealed a structure docked to the A-tubule between protofilaments A7 and A8 (Fig. 4E, Fig. 5A). In eel, this structure consists of two lobes connected to the DMT by a thicker proximal connection and thinner distal connection (Fig. 5B). The lobes of this structure reach out towards, and potentially contact, the B-tubule of DMT_n+1_. Like ODAs, this double-lobed structure maintains a 24-nm periodicity within the eel axoneme repeat unit (Fig. 5B)

In organisms that contain ODAs, the individual dynein motor domains are connected to the DMT through an ODA docking complex (ODA DC) (Gui et al., 2021a; Owa et al., 2014; Takada et al., 2002; Walton et al., 2021). These complexes bind to the DMT between the A7 and A8 protofilaments and each complex has at least two connection points. Single celled eukaryotes, such as *Chlamydomonas* and *Tetrahymena*, have a single-lobed ODA DC with one DMT contact directly below the lobe and another connection tailing away from the lobe towards the next most proximal ODA (Bazan et al., 2021; Gui et al., 2021a; Song et al., 2020). Multicellular animals, such as cows and sea urchins, tend to have double-lobed ODA DC structures and connections to the DMT are on either end of the lobes (Gui et al., 2021a; Lin and Nicastro, 2018). In fact, the structure of the bovine ODA DC (Gui et al., 2021a) resembles the density decorating the eel 96-nm repeat unit (Fig. 3G). Based on the location and periodicity of this structure, as well as its structural similarity to the bovine ODA DC, we predicted that the eel sperm flagellum contains an animal-like ODA DC complex.

The bovine ODA DC is made up of five proteins. The proximal attachment is a heterodimeric complex of CCDC151 and CCDC114. Calaxin (Efcab1) and ARMC4 constitute the proximal and distal lobes, respectively. TTC25 is the distal attachment between the ODA DC and the DMT (Gui et al., 2021a). Proteomic analysis of extracted eel flagella revealed three of these ODA DC-associated proteins, TTC25, ARMC4, and CCDC151 (Table 1). Calaxin and CCDC114 were not identified in the mass spectroscopy data, but the corresponding structures appear to be present in the eel ODA DC. It is currently unknown if these proteins are present but were not identified in the proteomic analysis or if these proteins have been replaced in the eel flagellum. The closely related European eel (*Anguilla anguilla*) encodes for predicted Ccdc114 and Efcab1 (calaxin) genes (GenBank).

To gauge the similarity of the eel and cow ODA DCs, we fit the cryo-EM structure of the bovine ODA DC (Gui et al., 2021a) into our eel sperm subtomogram average. The ARMC4, TTC25, and CCDC114/CCDC151 segments of the map fit within the cryo-ET density. The calaxin structure, on the other hand, does not fit into the map. In vertebrates, the calaxin subunit is situated between ARMC4 and the ODAs, facilitating ODA docking (Fig. 5E, (Yamaguchi et al., 2023)). While there does not appear to be density in the eel sperm average corresponding to calaxin, there is an as-yet unaccounted for density on the opposite side of the ARMC4 density. It is possible that in the eel axoneme, calaxin binds to the opposite side of the ARMC4 structure. Recently, it has been shown that calaxin is critical for ODA binding in zebrafish sperm flagella (Yamaguchi et al., 2023). In calaxin^-/-^ mutants, the ODAs, and the ODA-DC, are absent in ∼30% of selected particles. As calaxin is important for ODA docking, it is possible that an ancestral calaxin/ARMC4 mutation that forced calaxin to switch sides may be responsible for the loss of ODAs in the eel sperm flagellum.

### The inner dynein arms (IDAs) are not well-regulated in the eel sperm flagellum

As the eel sperm flagellum lacks ODAs, the IDAs are the only known protein motors within the axoneme that can drive ciliary beating. The 96-nm repeat average of the eel flagellum revealed a full complement of IDAs, including the six single-headed dyneins and the double-headed I1/*f* dynein (Fig. 4A). Like in other species, the eel IDAs sit below the DMT (towards the center of the axoneme) and reach out towards the neighboring DMT (Fig. 3E). Unlike other species (*e.g.,* sea urchin or *S. rosetta*), the eel IDAs are situated below the IJ of the neighboring DMT as opposed to behind the neighboring DMT (Fig. 3 E, see Discussion). IDAs utilize a stalk domain to reach out towards DMT_n+1_ (Lin et al., 2014). We searched for IDA stalks in both active and EHNA immobilized (Fig. 3) averages, but we did not find any corresponding density. Given the proximity of neighboring DMTs, one might expect features from the previous DMT (DMT_n-1_) to be present in the subtomogram average. While there is some weak density underneath the A-tubule, we were unable to determine if these were flexible/uncommon structures attached to the bottom of the A-tubule or if they were structures from the neighboring DMT.

In [9+2] flagella, several of the IDAs dock to the DMT through interactions with the RSs. To investigate how the eel IDAs dock onto the DMT, we compared eel sperm and *S. rosetta* 96-nm repeat averages at each of the IDA tails (Fig. 4E-T). The IDAs that do not normally interact with the RSs (IDAs *b*, *e*, and *g*) demonstrated a similar docking pattern between the two species. The docking of the remaining IDAs (*a*, *c*, *d*) is perhaps more interesting. Without the RSs, it would be expected that the docking would be radically different between the two species. Instead, it appears that the docking of IDAs *a*, *c*, and *d* is consistent between eel and *S. rosetta* (Fig. 4). IDA *d* typically docks to the DMT through an interaction with RS3, and this interaction appears to be retained using the eel RS3S complex. IDAs *a* and *c* dock via interactions with RS1 and RS2, respectively. As described above, there is density underneath the eel A-tubule that may correspond to FAP207 and FAP253. It is possible that these remnants of RS1 and RS2 are retained in the eel axoneme to facilitate docking of IDA *a* and IDA *c*.

In many axonemes of motile flagella, there is a set spacing of DMTs and a solid interaction between neighboring DMTs. For example, both the ODA and IDA binding sites are known for the sea urchin axoneme and neighboring DMTs interact in a defined manner, creating a regular register (Lin and Nicastro, 2018). To examine the inter-DMT register along the eel axoneme, we placed an isosurface rendering of the 96-nm repeat subunit back into the eel tomograms at the calculated positions using the “clonevolume” function in IMOD (Kremer et al., 1996). From these cloned volumes, we searched for a pattern of inter-DMT connections, focusing on the most prominent feature, the N-DRC. Analysis of these interactions revealed that not only does the inter-DMT register vary between flagella, but it can also vary between DMTs within the same flagellum. The lack of a specific inter-DMT register suggests that the ciliary beating of the eel flagellum may not be as well-regulated as that of other species, which may, in turn, lead to the helical beating pattern.

### The N-DRC is conserved in the eel flagellum

Although the eel flagellum lacks many of the canonical structural elements of the axoneme, a subset of these structures has been retained. One of the retained structures is the nexin-dynein regulatory complex (N-DRC). This complex binds to the bottom of the A-tubule and extends out to interact with the neighboring B-tubule (Fig. 6A). Many of the components of the N-DRC complex are visible in the subtomogram average of the eel axoneme repeat unit (Fig. 6 B). When viewed from underneath the DMT, the prongs of the baseplate scaffold (Fig. 6B) and both the proximal and distal lobes are visible (yellow arrowheads in Fig. 6). Proteomic analysis of extracted eel flagella revealed the presence of several N-DRC related proteins (Table 1). These proteins include DRC4 (baseplate/proximal lobe), DRC5 (L2 protrusion of the linker) and DRC11 (proximal lobe) (Gui et al., 2019).

**Figure 6:**
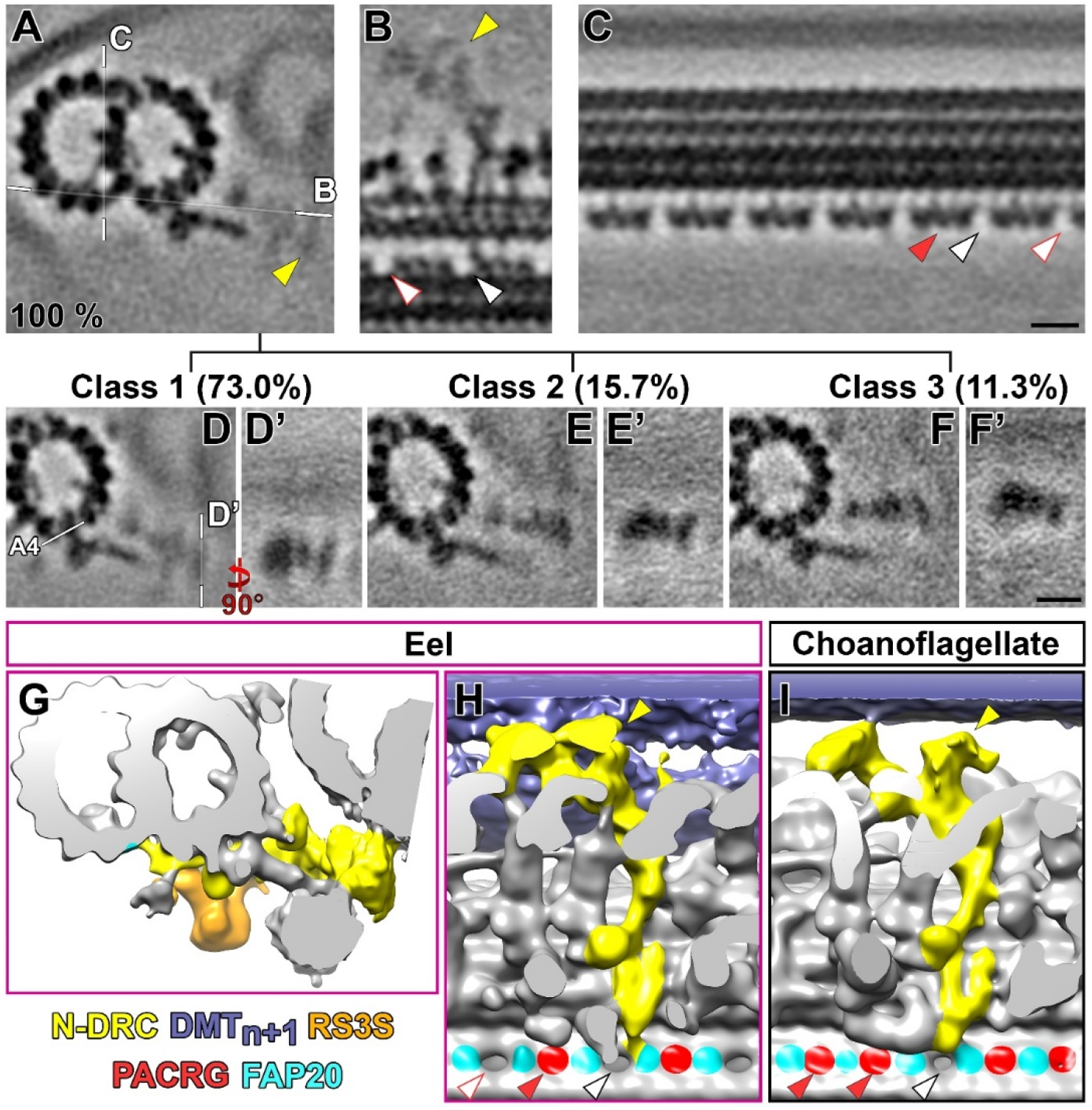
The eel sperm flagellum contains a flexible N-DRC and a unique inner junction (IJ) spacing. A-C) Cross sectional (A), bottom (B), and longitudinal (C) slices of the eel sperm 96-nm repeat average. The eel N-DRC (yellow arrowheads in A-B) extends from the bottom of the A-tubule towards the A-tubule of the neighboring DMT (A). There is strong density for the N-DRC baseplate (B), but the density is weaker for the proximal lobe and even weaker for the distal lobe, suggesting that the eel sperm N-DRC is flexible. Unlike most other known motile flagella, the eel sperm flagellum contains multiple holes along the inner junction (IJ). The black-outlined arrowhead points to the N-DRC-associated IJ hole and the red-outlined arrowhead points towards an additional hole. D-F) Class averages reveal structural flexibility in the eel sperm N-DRC. Cross sectional (D, E, F) and longitudinal (D’, E’, F’) slices of the eel average show the N-DRC in a low (class 1, D/D’), middle (class 2, E/E’), or high (class 3, F/F’) conformation. G-I) Isosurface renderings of the eel (G, I) and *S. rosetta* N-DRCs. In most organisms there is a single hole at the IJ located at the N-DRC (black-outlined arrowheads). The eel sperm flagellum has multiple holes along the length of the flagellum (red-outlined arrow), apparently due to a missing PACRG subunit (red arrowheads). Scale bars represent 10 nm.

Electron density corresponding to the proximal and distal lobes of the N-DRC was weaker than the surrounding structures in the eel axoneme subtomogram average (Fig. 6A-B). Weaker or blurred density in subtomogram averages can be caused by averaging flexible structures or by averaging together particles with and without the structure of interest. To investigate the cause of the blurred N-DRC densities, we performed classification on the N-DRC lobes. The classification revealed three different classes, with the main difference being the position of the N-DRC lobes (Fig. 6D-F). The most abundant class (73.0%) had the N-DRC extending from the A-tubule at a 20° angle (relative to protofilament A4). The other two classes, representing 15.7% and 11.3% of the particles, had A4-N-DRC angles of 10° and 2°, respectively (Fig. 6 E-F). Consequently, the N-DRC lobes were lower or higher, depending on the class, relative to DMT_n_. Class 1 (73.0%) had the lowest lobes and class 3 (11.3%) had the highest. The strength of the signal corresponding to DMT_n+1_ was also the strongest in class 1, suggesting that the interaction between neighboring DMTs may be stronger in this class.

### The inner junction of the eel flagellum demonstrates a unique protein pattern

The axonemal DMT is made up of a complete (13 protofilaments) A-tubule and an incomplete (10 protofilaments) B-tubule (Nicastro et al., 2011). There are two locations within the DMT where the A- and B-tubules interact. At the outer junction (the side of the DMT that faces the membrane), there is a unique series of interactions between two A-tubule protofilaments (A10/A11) and a B-tubule protofilament. On the opposite side of the DMT, the inter-tubule connections at the inner junction (IJ) are mediated by a series of non-tubulin proteins (Dymek et al., 2019; Khalifa et al., 2020; Khan et al., 2021). In most organisms, the IJ consists of a pair of peripheral proteins (in line with the tubulin protofilaments) and a series of MIPs located within the lumen of the B-tubule. The two peripheral IJ proteins are FAP20 and parkin co-regulated protein (PACRG) (Dymek et al., 2019). In animals, a third protein, meiosis expressed gene 1 product (MEIG1), is thought to bind to PACRG and the B-tubule protofilament along the outside of the microtubule (Khan et al., 2021; Li et al., 2015). Within the lumen of the B-tubule, there are a series of connections between tubules and between connecting proteins. Proteins involved in these connections include PACRG, MIP3a (FAP52), FAP276, FAP106, FAP126, and FAP252 (Dymek et al., 2019; Khalifa et al., 2020; Khan et al., 2021; Li et al., 2015). Some of these proteins, including FAP20, and PACRG, were found in the proteomic analysis (Table 1), and several others, including MEIG1, are encoded for in the *Anguilla anguilla* genome.

The periphery of the IJ consists of repeating subunits of FAP20 and PACRG (Dymek et al., 2019; Khan et al., 2021). In most organisms, this repetition is broken once every 96 nm at the N-DRC baseplate (Gui et al., 2019). In eel, however, there are multiple IJ holes present throughout the 96 nm repeat unit (Fig. 6C, H). These holes repeat at 16 nm intervals (Fig. 6C), suggesting that every other copy of either PACRG or FAP20 is missing from the structure. In *Chlamydomonas*, the IJ hole at the N-DRC is caused by a missing PACRG subunit (Gui et al., 2019). In the eel flagellum, there is a homologous hole at the base of the N-DRC and this hole fits into the periodicity of the other IJ holes (Fig. 6H). In contrast, the *S. rosetta* axoneme contains both PACRG subunits surrounding the main N-DRC IJ hole (Fig. 6I). Based on this continuity, we predict that the eel flagellum lacks every other copy of the PACRG protein. The periodicity of the IJ hole, along with the periodicity of the entire 96-nm repeat subunit, is controlled by the Ccdc39/Ccdc40 complex, also called the axonemal ruler (Oda et al., 2014). This complex is present in the eel flagellum, and there are no apparent structural differences between this complex in the eel flagellum and in other organisms. Thus, it is unlikely that the axonemal ruler is responsible for the eel’s unique IJ pattern.

Most organisms encode for a single isoform of PACRG. One notable exception is *Trypanosoma brucei*, that encodes for two PACRG proteins, called Tb PACRG A and Tb PACRG B (Dawe et al., 2005). As the eel flagellum has lost many of the structural elements of the axoneme, we wanted to determine if the eel had also lost a second isoform of PACRG. Using NCBI’s BLAST algorithm (Altschul et al., 1990), we searched for PACRG proteins across bony fish (*Actinopterygii*). We did not find any instances of multiple PACRG proteins, suggesting that the ancestral eel did not have a second copy that was lost over the course of its evolution. Even zebrafish (*Danio rerio*), recently shown to have a similar pattern of IJ holes as eel (Fig. S1E) (Yamaguchi et al., 2023), does not encode for two PACRG copies, suggesting that PACRG alone is not responsible for the unique IJ holes.

The canonical IJ hole at the N-DRC is caused by the baseplate inserting into the IJ, resulting in the loss of a PACRG subunit. To investigate the potential of another structure inserting into the IJ, we searched the eel flagellum for structures close to the IJ with a similar periodicity to the IJ hole. Several of the MIP structures displayed a similar periodicity as the IJ holes, including the MIP3 proteins that interact with the IJ itself. Each IJ hole sits at the position along the repeat unit where the MIP3a and MIP3b subunits connect (Fig. S1B). MIP3a is a relatively large MIP formed by the FAP52 protein, and MIP3b is the tether loop structure formed by FAP106 that both tethers FAP52 to the IJ and connects the A- and B-tubules (Khalifa et al., 2020; Song et al., 2020). The MIP3b structure spans ∼8 nm along the length of the DMT (Fig. S1), a distance that covers two FAP20 and one PACRG subunits. The IJ holes appear at the proximal end of the MIP3b structure and density from MIP3b protrudes slightly into the hole. Published DMT averages from other organisms, including mice, *Chlamydomonas* (Fig. S1F), *Tetrahymena*, and cows (Gui et al., 2021a; Lin et al., 2019; Song et al., 2020), display MIP3b density in this region interacting with the PACRG subunit. The only other organisms that have been reported to have additional IJ holes include *S. rosetta* (there is a DMT-specific loss of another PACRG subunit, (Pinskey et al., 2022)) and zebrafish (Yamaguchi et al., 2023) with a similar pattern as eel. At the canonical IJ holes (at the N-DRC), the MIP3b of these organisms also protrude slightly into the IJ hole. These protrusions make it unlikely that there is extra density in the eel FAP52 protein that causes the loss of a PACRG subunit.

### The eel flagellum contains a rail-like microtubule inner protein (MIP) and a unique outer junction complex

Subtomogram averaging of the eel sperm flagellum revealed that there is a rail-like structure within the lumen of the A-tubule (Fig. 7A, G-I). This structure is similar to the rail-MIP seen in *S. rosetta* (Pinskey et al., 2022) or the mouse MIP4 (Gui et al., 2021a) and is not present in the axonemes of most species studied to date. The rail-MIP runs the length of the 96-nm repeat unit and spans between protofilaments A12 and A1 (Fig. 7G). There are apparent connections between the rail-MIP and protofilaments A1 and A12, but not A13 between the two (Fig. 7H-I). The function of these Rail-MIPs remains unknown, but we hypothesize that they could play a similar role as tektins do in several mammalian axonemes (Gui et al., 2021a; Song et al., 2020).

**Figure 7:**
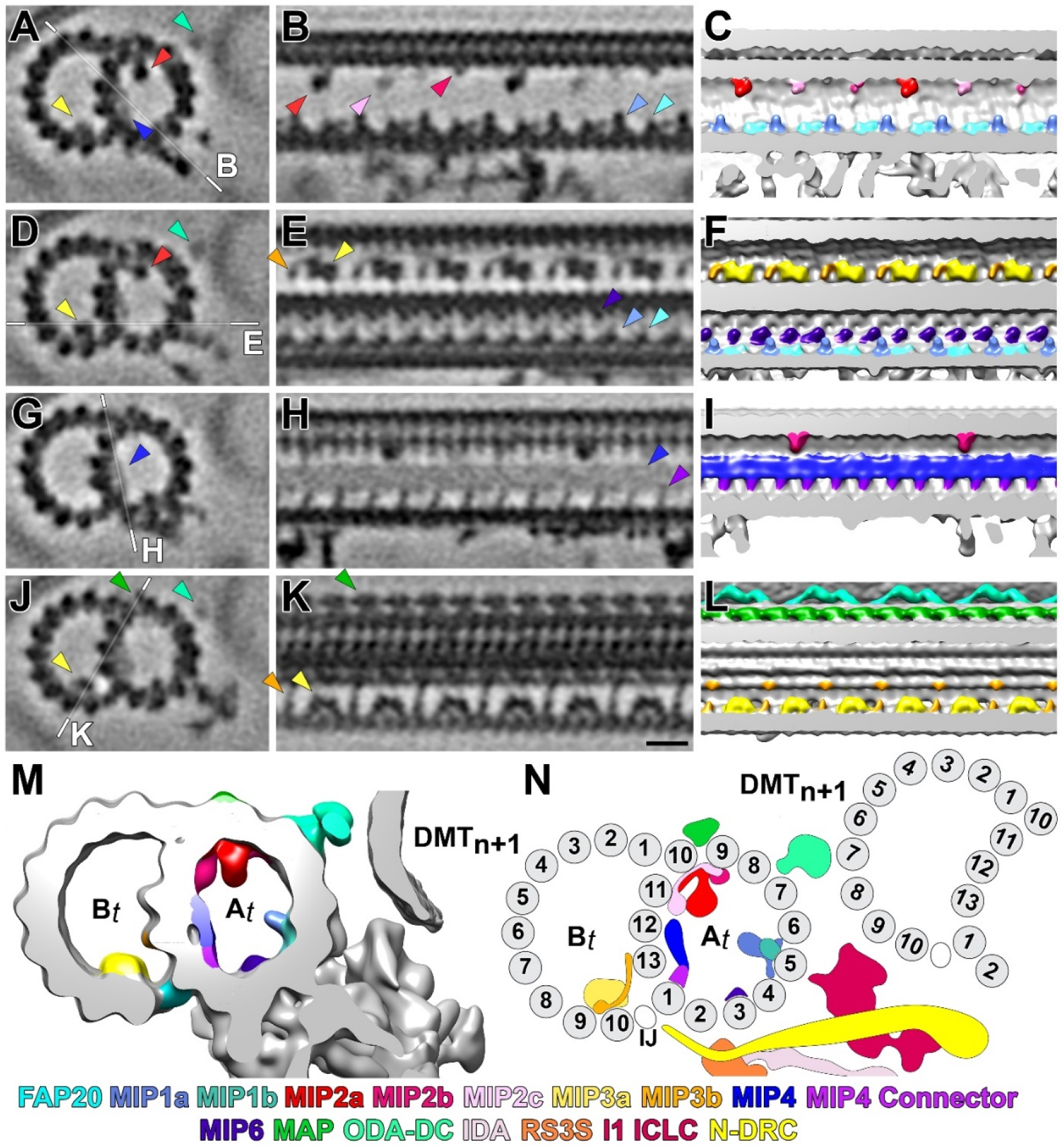
The eel flagellum contains both conserved and unique microtubule-associated proteins. A-L) Cross-sectional (A, D, G, L) and longitudinal (B, E, H, K) slices of the eel sperm average and the corresponding isosurface renderings (C, F, I, L). MIPs appear to be conserved in the eel sperm flagellum (MIP1, MIP2, MIP3, MIP4, MIP6). Colored arrowheads highlight specific MIPs and are colored according to the key. The eel MIP4 is similar to the so-called Rail-MIP seen in *S. rosetta* and the mouse MIP4 but appears to have discrete connections with the A-tubule (purple). The “outer junction” of the eel sperm DMT contains a unique microtubule accessory protein (MAP, green) that spans the gap between protofilaments A10 and B1. White lines in the cross sections represent the location of the subsequent images. M) Isosurface rendering of the eel sperm 96-nm repeat average. Individual microtubule-associated proteins (MIPs, ODA-DC, etc.) are colored according to the key. N) Cartoon representation of the eel sperm MIP-decorated eel sperm DMT. The scale bar in K represents 10 nm and is valid for all tomographic slices.

Much like the N-DRC lobes, the electron density of the rail-MIP in the full subtomogram average was weaker than the surrounding protofilament density (Fig. 2E). To determine the cause of the blurred density, we performed classification on the rail-MIP structure. The classification yielded two classes, one with the rail-MIP (51.8% of particles) and one without the rail-MIP (44.3% of particles). Many species have DMT-specific (*e.g.,* the 5-6 bridge in *Strongylocentrotus*) or proximal-distal specific (*e.g.,* the 1-2 bridge in *Chlamydomonas*) structures within their axonemes (Lin et al., 2012). To investigate possible proximal-distal relationship or DMT-specificity, we mapped the classed particles back onto the individual flagella. The presence or absence of the rail-MIP appeared to be sporadic throughout the flagella. It does appear, however, that compressed flagella tended to have more particles that did not contain the rail-MIP. It is currently unknown if the relative absence of the rail-MIP is a symptom or a cause of the increased flagellar compression.

Alongside the rail-MIP, the structure of the eel 96-nm axoneme repeat unit contains several unique structural features. The first of these features is an apparent microtubule accessory protein (MAP) that connects protofilaments A9 and A10 (Fig. 7K-M). MAPs are proteins that attach to the exterior of the microtubule doublets, and they are thought to play roles in DMT stability and flagellar signaling (Linck et al., 2014; Ma et al., 2019; Nicastro et al., 2011). A similar, albeit smaller, bridging structure has been identified spanning A9-A10 in *Chlamydomonas* (Ma et al., 2019), but the protein identity of this structure is unknown.

## Discussion

### The eel flagellum employs a unique inter-DMT connection strategy to propel ciliary beating

In traditional axonemes, binding of the ODAs, IDAs, and the N-DRC is largely conserved (Gui et al., 2019; Lin and Nicastro, 2014; Lin and Nicastro, 2018). These structures dock to the A-tubule of DMT_n_ and interact with the B-tubule of DMT_n+1_. The eel axoneme, on the other hand, does not adhere to these canonical inter-DMT interactions (Fig. 1, Fig. 3). For example, the distance between neighboring DMTs in the eel sperm flagella (∼ 40 nm on average) is smaller than that seen in [9+2] flagella (*e.g.,* ∼70 nm in sea urchin (Lin and Nicastro, 2018)). To better illustrate the differences in inter-DMT interactions between [9+2] flagella and the [9+0] eel flagella, we modeled the protofilaments and inter-DMT connections from the eel subtomogram average and from the *S. rosetta* axoneme repeat unit (Fig. 8). The protofilaments of DMT_n_ were manually annotated and then were copied for estimation of DMT_n+1_. Alignment and protofilament assignment for DMT_n+1_ was estimated by matching the B-tubule-to-A-tubule angle of the neighboring DMT with special care taken to align protofilaments A10-A1 and B1/B10.

**Figure 8:**
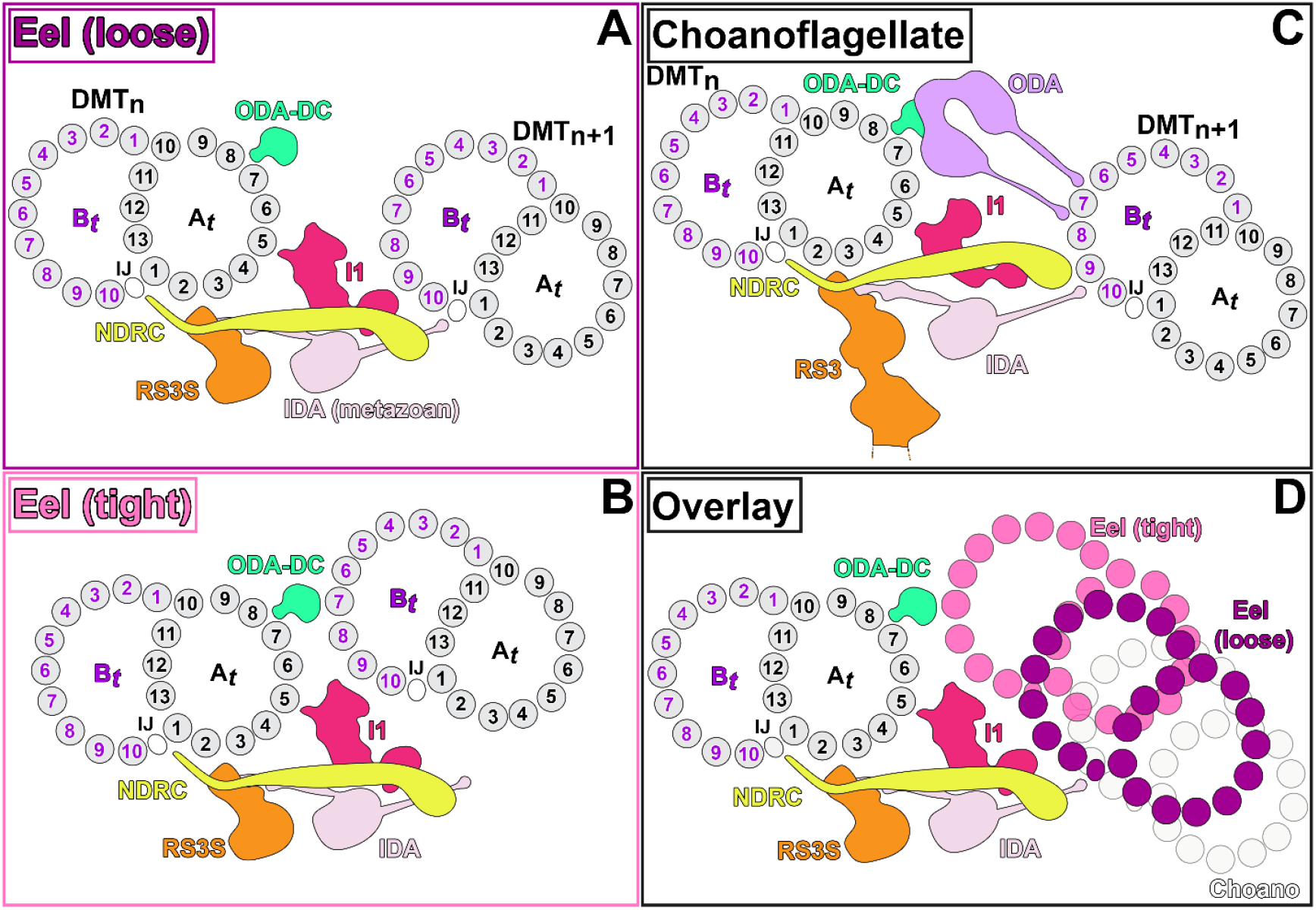
Comparison of the inter-DMT connections of the [9+0] (eel) and [9+2] (*S. rosetta*) axonemes. A-B) Cartoon representations of the inter-DMT connections (IDAs, I1 ICLC, N-DRC, ODA-DC) present along the eel flagellum. IDA stalks were not visible in the eel sperm tomograms or averages, so the typical metazoan IDA orientation is displayed for reference. The eel sperm inter-DMT spacing is variable between and within flagella. The “loose” conformation (A) positions the IDA near the neighboring DMT, but it is unclear where, or if, the IDA binds in the “tight” conformation (B). C) Cartoon representation of the inter-DMT interaction present in the choanoflagellate axoneme. The choanoflagellate RS3 extends further towards the CPC and has been truncated in this representation. D) Overlay of the three inter-DMT connections (tight eel, loose eel, choanoflagellate). Note the spatial difference between the two eel states and the similarity of the loose eel conformation to the typical metazoan spacing. For all cartoons, the DMT-to-DMT spacing was established by manual assignment of protofilaments in DMT_n_ and estimation of the A-tubule B-tubule angle in DMT_n+1_.

In the *S. rosetta* axoneme, the ODAs, IDAs, and the N-DRC all bind to the B-tubule of the neighboring DMT (Lin and Nicastro, 2018). The ODA stalks bind to DMT_n+1_ in the grooves between protofilaments B6, B7, and B8 (Fig. 8). The IDA stalks dock into the groove between B9 and B10 in the neighboring DMT, and the N-DRC interacts with B9. The eel flagellum lacks ODAs, so it necessarily lacks any interactions between ODA stalk subunits and the neighboring DMT. The lack of ODAs does not mean that the eel flagellum lacks any outer connections with the neighboring DMT; the eel ODA DC extends out towards DMT_n+1_ and appears to connect to the B-tubule (Fig. 5B & D, Fig. 7A). Based on the location of the protofilaments of DMT_n+1_, we predict that the eel ODA DC contacts the neighboring doublet around B7, the middle protofilament involved in true ODA binding.

The interactions between the IDA stalk domains and the neighboring DMT are not directly visible in the current subtomogram averages. The IDA dynein heads themselves, however, sit below the inner junction region of the neighboring doublet (Fig. 3 E, Fig. 4). Based on this positioning, it unlikely that the eel IDAs interact with the neighboring B-tubule. If the IDA stalks were to bind between B9-B10, they would have to be positioned vertically with respect to the dynein motor, if not slightly further back towards DMT_n_. Although it is not impossible, this orientation goes against much of what is known about the structure and function of dynein motors (King, 2016; Lin and Nicastro, 2014; Lin and Nicastro, 2018). More likely, the IDA stalks of the eel flagellum bind to the A-tubule of the neighboring DMT.

The N-DRC of the eel flagellum presents another interesting puzzle. The N-DRC appears to reach past the neighboring B-tubule and, much like the IDAs, is positioned around the inner junction of the neighboring doublet (Fig. 6A). In the two most abundant classes from the N-DRC classification (Fig. 6D-E), the N-DRC lobes do not appear to be pointing towards the neighboring protofilaments. In fact, it appears that the N-DRC is pointing towards a structure that is hanging off the bottom of the A-tubule. Initially, we hypothesized that the N-DRC was interacting with the RS3S structure, the only structure that hangs below the eel A-tubule. To investigate the inter-doublet register in the eel flagellum, we placed an isosurface rendering of the subtomogram average back into model points used for particle alignment (clonevolume feature in PEET). Using this model, we searched for a pattern in the inter-DMT connections along the flagellum. Unfortunately, the register appeared to be random, and we were unable to identify a specific binding pattern. Without ODAs, the eel flagellum lacks 12 (or 8) inter-DMT connections per axoneme repeat. Even though the ODAs are not permanently attached to the neighboring doublet (Lin and Nicastro, 2018), it may be that these additional connections assist in creating the proper inter-DMT register in [9+2] flagella.

### The helical beating pattern of the eel flagellum is likely caused by the lack of signal transduction machinery

Motile cilia beating is controlled through coordinated regulation of dynein motor activity (Lin and Nicastro, 2018). Although the precise mechanism(s) of signal transduction remain(s) elusive, the CPC, RSs, and both rows of dynein arms are known to play a role (Adams et al., 1981; Diener et al., 1993; Lin and Nicastro, 2018; Viswanadha et al., 2017; Wirschell et al., 2011). When these structures are missing or defective, flagellar beating is abnormal or paralyzed. For example, the CPC-less *pf18* mutant in *Chlamydomonas* has completely immotile flagella (Adams et al., 1981) and the RS-less *pf14* mutant has paralyzed flagella (Diener et al., 1993). Loss of individual proteins or complexes can also lead to atypical flagellar beating patterns. For example, when CPC1, an adenylate kinase protein in the CPC, is knocked out, the flagella have an altered beating frequency (Zhang and Mitchell, 2004). Similarly, the *pf17 Chlamydomonas* mutant lacks radial spoke heads, but retains the stalks, and has altered flagellar beating that causes the cells to wiggle instead of swim (Huang et al., 1981). *Chlamydomonas* mutants that lack ODAs and IDAs display slowed flagellar beating (Kamiya et al., 1991; Perrone et al., 1998; Rupp et al., 1996; Sakakibara et al., 1991).

The eel sperm flagellum does, in fact, beat, albeit with a helical pattern (Movie S1) (Woolley, 1998b). While the eel flagellum doesn’t beat in the quasi-planar manner of [9+2] flagella, it does appear to have a regular beating pattern (Movie S1) (Woolley, 1997; Woolley, 1998b). In Movie S1, there are many cells that appear to be immotile, but there are also several cells that are actively swimming. Whereas the beat pattern and frequency qualitatively differ from cell to cell, they all exhibit a beating pattern, suggesting some form of dynein regulation must be taking place. Without regulation, all the dyneins would fire at the same time, generating a net zero force on the flagellum and not initiating bending (Lin and Nicastro, 2014). The mechanism behind the regulation of eel flagellum dynein motors remains unknown. As the eel flagellum lacks the CPC and RSs, key pieces of signal transduction in other organisms, the regulation of dynein activity must come from either the IDAs themselves or from an as-yet undetermined signaling factor.

### Evolutionary pressures may have led to the “minimum ensemble” required for flagellar beating

The canonical [9+2] axoneme is nearly universal amongst sperm flagella (Satir and Christensen, 2007). The eel sperm flagellum, on the other hand, has lost many of the structures that are known to drive and regulate ciliary beating. Although it lacks these critical structures, the eel flagellum still beats (Movie S1, (Woolley, 1997)) and this beating is sufficient to facilitate reproduction (van Ginneken and Maes, 2005). As the eel flagellum still beats and is sufficient for reproduction, the eel flagellum appears to contain a minimum ensemble of axonemal proteins required for life.

The question then arises, what evolutionary factors lead to the structural cassette of the eel axoneme? Organisms as far apart evolutionarily as *Chlamydomonas*, choanoflagellates, cows, and humans share a common set of flagellar components (Coutton et al., 2018; Gui et al., 2021a; Nicastro et al., 2011; Pinskey et al., 2022). The eel flagellum likely diverged from a common ciliated ancestor and lost structural features over time. For example, the presence of an ODA-DC, but not full ODAs, suggests that the full feature was present in the axoneme at some evolutionary timepoint. Similarly, the RS3S structure and remnants of RS1 and RS2 in the eel flagellum suggests that some ancestral eel sperm flagellum may have contained full RSs.

If the American eel did lose its axonemal components over time, there must have been an evolutionary advantage, or at least no evolutionary penalty, for doing so. Atlantic *Anguillid* eels travel to a small region in the Sargasso Sea to spawn (Jacobsen et al., 2014b; Munk et al., 2010; Pujolar et al., 2014). These migrations can cover thousands of kilometers; ∼4,000 km on average for the American eel and 5,000-10,000 km for the European eel (Beguer-Pon et al., 2015; Mccleave and Kleckner, 1982). Pacific eels (*Anguilla* japonica) endure similarly long migrations in the hopes of passing along their genetic material (Tsukamoto et al., 2011). The eels are thought to swim constantly throughout these migrations and are not thought to feed during the entire journey (van Ginneken et al., 2005a; van Ginneken and Maes, 2005). Much like migratory salmon (von Schalburg et al., 2018), these eels undergo significant morphological changes during their migration. Although spawning American or European eels have not been caught in the wild, Japanese eels have been (Tsukamoto et al., 2011; van Ginneken and Maes, 2005). Once the eels reach the open sea during their migration, digestive tract begins to atrophy and the gut is lost (Tsukamoto et al., 2011). Without feeding, the eels must rely on their energy stores to not only travel thousands of kilometers, but also develop gonads and produce gametes (e.g., sperm cells).

The traditional [9+2] axoneme consists of over 600 unique proteins (Nicastro et al., 2006). With an average length of ∼35 µm (Woolley, 1997), there are ∼3300 96-nm repeat units along the length of the eel flagellum. Dynein heavy chains are approximately 4000 amino acids (Uniprot), so by simply removing the ODAs from the axoneme, the eel no longer needs to synthesize and assemble 5.26 × 10^7^ amino acids per flagellum. Coupling in the lack of full RSs and the CPC, the eel axoneme represents a significant energy savings compared to traditional [9+2] axonemes. By spending less energy per sperm cell, male eels could either devote more energy to migration (increasing the likelihood they survive to reproduce) or generate more sperm cells (increasing their likelihood of passing along their genetic material during mating events).

In captivity, *Anguilla* can release > 10 × 10^9^ sperm cells per mL. This is a higher sperm density than most terrestrial animals, including bulls (∼1 × 10^9^ sperm/mL), boars (∼3 × 10^8^ sperm/mL), rams (∼3 × 10^9^ sperm/mL), and humans (∼4 × 10^7^ sperm/mL) (Sengupta et al., 2018; Setchell, 1997). This concentration is in line with other aquatic species, including rainbow trout (∼12 × 10^9^ sperm/mL), yellow perch (∼41.5 × 10^9^ sperm/mL), and whitefish (∼8 × 10^9^ sperm/mL) (Ciereszko and Dabrowski, 1993). This is also similar to other marine fish species such as turbot (∼2 × 10^9^ sperm/mL), tuna (∼40 × 10^9^ sperm/mL), sea bass (∼30 × 10^9^ sperm/mL), and cod (∼6.5 × 10^9^ sperm/mL) (Cosson et al., 2008) that have retained the [9+2] axoneme. Salmon, another group of catadromous fish, also undergo dramatic physiological changes during their breeding migrations (von Schalburg et al., 2018). Unlike eels, however, salmon still utilize a [9+2] axoneme in their sperm flagella. This retention within another energy starved fish species suggests energetics alone may not account for the [9+0] nature of the eel axoneme.

Although the exact mating behaviors of eels remain a mystery, there is some information available about their breeding habits (Beguer-Pon et al., 2015; Guarniero et al., 2020; Jacobsen et al., 2014a; van Ginneken and Maes, 2005). Eels, like many fishes, release both their eggs and sperm into the ocean for fertilization (Guarniero et al., 2020; van Ginneken and Maes, 2005). Unlike salmon, that form mating pairs and deposit their sperm onto already placed eggs (von Schalburg et al., 2018), eels are thought to engage in mass spawning events. Captive eels have been shown to engage in “mass spawning” events when multiple males spawn in response to hormonal triggers (Okamura et al., 2014; van Ginneken et al., 2005b). In fact, male eels have been observed swarming around released eggs and releasing their milt (Okamura et al., 2014). Through a combination of communal breeding and “mass spawning,” it is likely that individual eel sperm cells do not need to migrate long distances to fertilize the egg. The relatively short (1-10 minute) beating time of the activated eel sperm flagellum (Movie S1, (van Ginneken et al., 2005b; Woolley, 1998a; Woolley, 1998b)) provides evidence in support of this hypothesis. If this hypothesis holds true, it is likely that the loss of prolonged and coordinated flagellar beating would pose little to no evolutionary disadvantage for the eel. Indeed, even a minimal competitive disadvantage from the flagellar beating would likely be outweighed by energetic advantages. Thus, we hypothesize that the eel was able to lose its ODAs, RSs, an CPC due to the evolutionary advantage conferred by the energy saved from making a minimal axoneme.

An alternative explanation for the loss of numerous structures from the eel sperm axoneme lies in the composition of the eel oocyte. As described above, the eel sperm does not need to travel large distances to reach a suitable oocyte. Over long distances, the regulated, sinusoidal beating of the traditional [9+2] axoneme is more energy efficient and generates more forward thrust than the eel’s helical beating pattern (Woolley, 1998a). The helical beating pattern does, however, generate more torque at the tip of the sperm head than traditional planar beating (Woolley, 1998b). This torque may be beneficial as the sperm cell attempts to penetrate the egg. Mammals, including humans, alter the beating of their sperm flagella during the final stage of fertilization (Austin, 1951; Chang, 1951; Zhao et al., 2022). This process, called capacitation, results in increased beat frequency at the cost of beat accuracy (inconsistent beat amplitudes). It is possible that during mass spawning the “pre-capacitated” eel sperm cell is at an advantage compared to a sperm cell with a traditional, planar beat pattern.

Regardless of the evolutionary rationale, the tail of the eel sperm cell contains one of the few known examples of a non-[9+2] motile cilium (Buceta et al., 2005; Schrevel and Besse, 1975; Woolley, 1997). As shown previously (Woolley, 1997) and characterized in this manuscript, the eel axoneme lacks many of the structures that are known to drive and regulate ciliary beating. Even without these structures, the eel flagellum still beats (Movie S1, (Woolley, 1997)), suggesting that the structures present in the eel axoneme are sufficient for ciliary beating. The ODAs, RSs, and the CPC, all structures known to be essential for ciliary beating in other organisms, are missing from the eel flagellum. The structures present in the eel sperm flagellum represent a “minimum ensemble” of proteins required for ciliary beating. Investigations of the signaling pathways and structural changes that facilitate eel sperm flagellar beating will provide information about the basic phenomena that drive ciliary beating.

## Materials and Methods

### Eel Maturation, Sperm Isolation, and Motility Movies

Sexual maturity was artificially triggered in captive wild-caught eels through hormone injections as described in (Oliveira and Hable, 2010). Briefly, eels were injected with Human Chorionic Gonadotropin (HCG) hormone weekly. Spermatogenesis was checked by hand stripping. Spermatozoa were extracted from mature male eels through strip-spawning (Koumpiadis et al., 2021). Approximately two mL of sperm sample was collected from each of four mature male eels. For immobilized samples, extracted sperm cells were treated with a final concentration of 2 mM EHNA (erythron-9-(2-hydroxy-3-nonyl)adenine, Sigma) and incubated for five minutes at room temperature. For live cell fluorescence imaging, cells were treated with a final concentration of 50 µg/mL of FM1-43 dye and incubated in the dark for one minute at room temperature. Sperm cell motility was activated by diluting the sample 1:1000 in artificial sea water (360 mM NaCl, 50 mM MgCl_2_, 10 mM CaCl_2_, 10 mM KCl, and 30 mM HEPES, pH 8.0). The eel sperm began to lose activity within minutes of activation, so movies were collected, or the samples were plunge frozen, within 30 seconds of dilution in sea water. Movies of actively swimming sperm cells were collected at the University of California, Santa Barbara using a Nikon Ti-2 microscope and an Andor-Zyla camera running Micro-Manager 1.4.23.

### Grid Preparation and Cryo-Electron Tomography

Cryogenic samples were prepared as described previously (Lin and Nicastro, 2018). Briefly, holey carbon grids (Quantifoil R2/2) were glow discharged for 30 seconds in a Pelco EasyGlow glow discharge unit. Eel sperm samples were activated with sea water (see above) and quickly mixed with BSA-coated colloidal gold nanoparticles to be used as fiducial markers. The grids were loaded onto a homemade manual plunge freezing device and loaded with 3-5 µL of sample. Excess sample was blotted away by gently applying filter paper (Whatman #1) to the back side of the EM grid. The grids were then immediately plunged into liquid ethane, maintained at -175 °C. Frozen grids were stored in liquid nitrogen until they were examined by TEM.

Vitrified grids were transferred into a grid clipping station (Thermo Fisher Scientific) and clipped into Autogrid rings. The clipped autogrids were then transferred into a Titan Krios TEM (Thermo Fisher Scientific) for tilt series collection. Tilt series were collected from -60° to +60°, with images collected every 2° using a dose-symmetric collection scheme (Hagen et al., 2017) controlled by SerialEM (Mastronarde, 2005). Images were recorded using a 5k × 4k K3 Summit direct electron detector (Gatan) in counting mode (20 frames per second, 0.75 second exposures, 1.6 electrons/Å^2^ per second). A nominal magnification of 26,000x was used for data collection with an effective pixel size of 3.15 Å. Images were collected using a Volta-Phase-Plate (Danev et al., 2014), operated at -0.5 µm defocus, and a post-column energy filter (Gatan), operated in zero-loss mode with a 20-eV slit width. The total electron dose per tilt series was limited to <110 e^-^/ Å^2^.

### Image Processing and Subtomogram Averaging

Movie frames were dose-weighted and motion corrected using MotionCor2 (Zheng et al., 2017). Tilt series preprocessing and tomogram reconstruction was performed using the IMOD software package (Kremer et al., 1996). The 10-nm gold nanoparticles added to the sample prior to vitrification were used as fiducial markers. 3D reconstructions were calculated using weighted back-projection. Tomograms with compressed axonemes or evidence of crystalline (non-vitreous) ice were excluded from subsequent analyses.

Subtomogram averaging was performed using the PEET package (Nicastro et al., 2006). Individual 96-nm axoneme repeats were picked from each of the nine DMTs within the eel flagella. The traditional axoneme subtomogram averaging workflow in the Nicastro lab places the initial model points for particle picking between RS1 and RS2 (Lin and Nicastro, 2018). The eel flagellum lacks RSs, and the only prominent features decorating the DMTs are the I1 ICLC and the N-DRC, but these structures were not readily visible in the raw tomograms. To ensure the particles along each DMT were in the same register, we first performed individual PEET runs for each DMT by itself. The I1 ICLC was visible in the individual DMT averages, and we centered each particle at the bottom of the A-tubule below the base of the ICLC. Model points were moved into the proper register using an in-house Matlab-based special feature retrieval program. Properly centered points were then aligned and averaged using PEET. Classification was performed using the unsupervised classification protocol built into the PEET program (Heumann et al., 2011). Soft-edge masks were used to select regions of interest for classification (*i.e.,* the rail-MIP or the N-DRC lobes). Isosurface renderings of the subtomogram averages were generated using UCSF Chimera (Pettersen et al., 2004) or ChimeraX (Pettersen et al., 2021). Tomographic slices were denoised using a weighted median filter (the smoothing filter in IMOD) to improve the contrast for presentation. Subtomogram averages were calculated from raw tomograms. Additional denoising of tomograms collected from thicker regions of ice was conducted using cryoCARE (Buchholz et al., 2019). Briefly, even- and odd-numbered frames were reconstructed separately. 1200 subvolumes (64 voxels) were extracted and used to train the neuronal network (batch size 16, learning rate 0.0004, 200 epochs, 75 training steps per epoch) and the trained network was then applied to the full (odd and even frames) reconstruction for denoising. Prior to training, a mask was applied around the flagellum to focus the denoising.

### Proteomic Analysis

Eel sperm flagella were separated from the sperm heads as described previously (Woolley, 1997) with minor modifications. Briefly, isolation buffer (250 mM NaCl, 1 mM MgSO_4_, 0.1 mM EGTA, 2 mM KCl, 10 mM Tris-HCl (pH = 8.8), 20 mM glucose) was added to suspended sperm cells and the reaction was mixed on a rotary mixer for 10 minutes at 4 °C. Cells were then transferred to ice-cold microcentrifuge tubes and vortexed for 10-15 seconds (∼3000 revolutions/minute) to separate flagella from sperm heads. Flagella detachment was monitored via light microscopy. Detached sperm heads were separated via centrifugation (5 min. at 1500 rpm, 4 °C) and the flagella-containing supernatant was pelleted using 10,000 × g for 10 min. Isolated flagella were resuspended and stored in HMEEK buffer (30 mM HEPES, 25 mM KCl, 5 mM MgSO_4_, 0.1 mM EDTA, 1 mM EGTA, pH 7.2) and stored at -20 °C.

Flagellar proteins were separated via SDS-PAGE using a 4-12% gradient SDS-polyacrylamide gel. The gel was stained via Coomassie brilliant blue and then destained to minimize background signal. Sample-containing gel lanes were excised from the gel and were cut into ∼1 mm^3^ pieces. In-gel trypsinization and MS peptide identification were performed by the University of Texas Southwestern Medical Center Proteomics Core Facility. Identified peptide fragments were compared against the *Anguilla rostrata* (American eel), *Anguilla anguilla* (European eel), *Danio rerio* (zebrafish), *Mus musculus* (mouse), and human proteomes using the CPFC (Trudgian et al., 2010) or Proteomics Discoverer (Thermo Fisher Scientific) programs. Known axonemal proteins are reported in Table 1, although proteins with < 5% coverage and fewer than 3 peptide spectral matches (PSMs) were excluded. Dynein proteins were labeled according to the unified dynein nomenclature laid out in (Braschi et al., 2022).

## Acknowledgements

The authors would also like to thank Dr. Daniel Stoddard for providing management and training at the UT Southwestern Cryo-Electron Microscopy Facility. Computational resources were provided, in part, by the UT Southwestern Medical Center BioHPC computing cluster located in the Lydia Hill Department of Bioinformatics. Mass spectroscopy was carried out in the UT Southwestern Proteomics Core Facility. Cryo-ET data were collected in the UT Southwestern Medical Center Cryo-EM Facility, which is supported in part by the Cancer Prevention and Research Institute of Texas (CPRIT) Core Facility Support Award RP170644. These experiments were funded by the following grants: R01GM083122 (to D.N.) and a CPRIT grant (RR140082) to D.N. J.R.S. is a Simons Foundation awardee of the Life Sciences Research Foundation.

## Figures/Figure Legends

**Figure S1:**
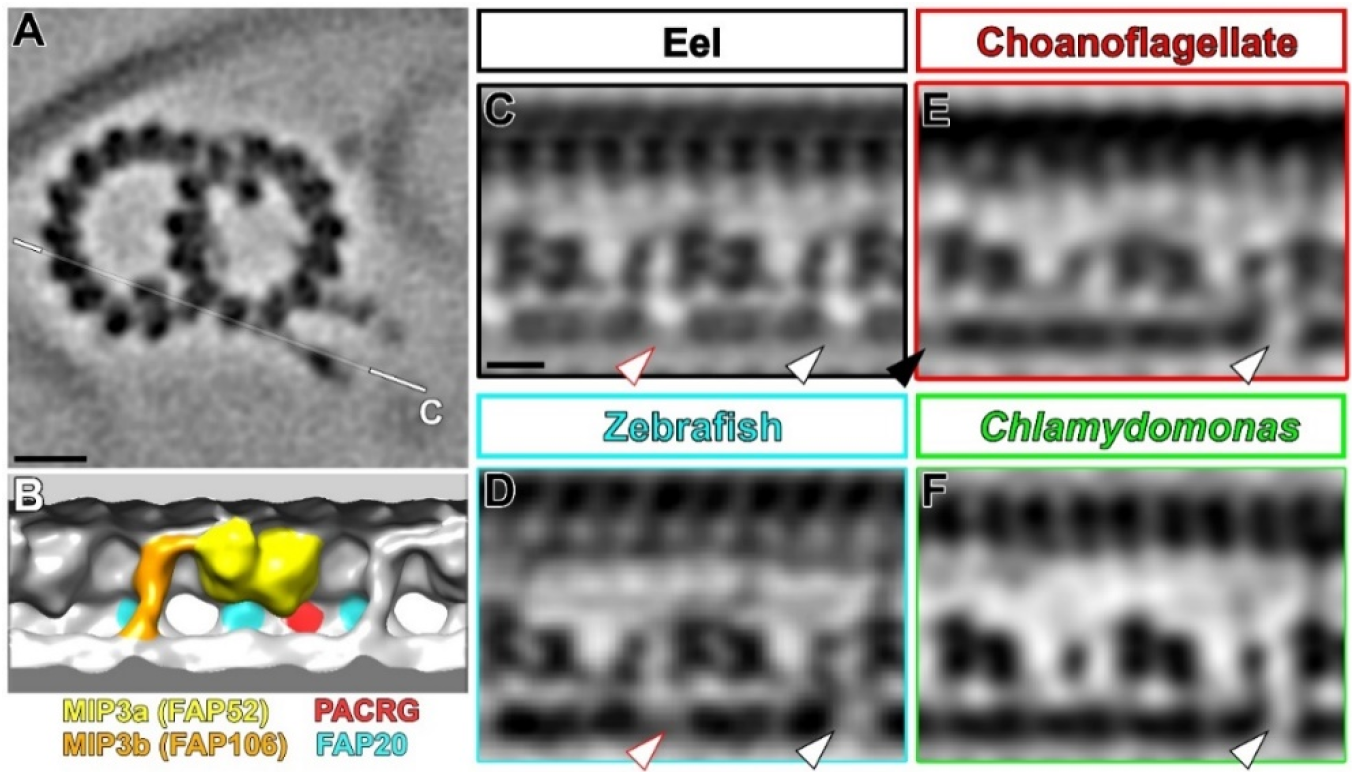
MIP3 (FAP52) does not extend into the IJ holes along the eel sperm flagellum. A) Cross-sectional slice of the eel 96-nm repeat average displaying an IJ hole and a MIP3b subunit. The white line indicates the position and orientation of the slice in C. B) Isosurface rendering of MIP3 and the IJ. Note the absence of any protein density extending into or near the hole, suggesting that it is not allosteric blockage that leads to the missing PACRG subunits. Structures in B are represented by the colors depicted in the key below B. C-E) Tomographic slices of the eel (C), zebrafish (D, EMD 34791), *S. rosetta* (E), and *Chlamydomonas* (F) MIP3-IJ regions, depicted in the same orientation as B. The distal lobe of MIP3A lies adjacent to the IJ hole but does not appear to extend into the hole. The main IJ hole (black outlined arrowheads) is aligned in each slice and is the only hole present in the *Chlamydomonas* average. The *S. rosetta* contains weaker density at a second PACRG site (black arrowhead) that depicts a second, DMT-specific IJ hole. The zebrafish sperm axoneme appears to share a secondary PACRG hole (red outlined arrowhead) with the eel sperm flagellum. Scale bars represent 10 nm in A and 5 nm in C (valid for C-F).

## Notes

### Competing Interest Statement

The authors have declared no competing interest.

